# Cyanobacteria using urea as a nitrogen source can overcome acid stress

**DOI:** 10.1101/2023.03.29.534730

**Authors:** Shuang-Qing Li, Hai-Long Huang, Tao-Ran Sun, Hua-Yang Gao, Xin-Wei Wang, Fei-Xue Fu, David A. Hutchins, Hai-Bo Jiang

**Affiliations:** Key Laboratory of Marine Biotechnology of Zhejiang Province, School of Marine Sciences, Ningbo University, Ningbo, Zhejiang 315211, China; Southern Marine Science and Engineering Guangdong Laboratory (Zhuhai), Zhuhai, Guangdong, 519080, China; Department of Biological Sciences, University of Southern California, Los Angeles, CA 90089, U.S.A.

**Keywords:** Cyanobacteria, Acid stress, Urea utilization, Urease, *Helicobacter pylori*

## Abstract

Cyanobacteria play essential roles in marine primary productivity and the global carbon/nitrogen biogeochemical cycle. Increasing urea emissions and decreasing pH value in the ocean caused by human activities are changing the community structure and competitive interactions of marine phytoplankton, which will have a profound impact on the marine ecosystem and global biogeochemical cycle. Here, we report that a coastal *Synechococcus* strain exhibited better adaptability to extreme low pH conditions when it uses urea as nitrogen source compared to using other inorganic nitrogen. Very low pH values can also alleviate damage by high concentrations of urea to cyanobacteria. Urease plays an essential role in this process. *Synechococcus* mutants with inactivated urease cannot adapt well to highly acidic environments, while heterologous expression of urease homologs from acidophilic *Helicobacter pylori* can help the cyanobacterial mutants to restore their adaptability to acidification. A TARA Oceans database analysis indicates that the distribution of cyanobacteria with the urease gene is closely related to estuaries and nearshore waters with potentially high urea inputs. In summary, we report for the first time that the use of urea and adaptation to acid stress are highly interactive in marine phytoplankton. Future work should determine whether this interaction is likely to allow phytoplankton that utilize urea to have a competitive advantage in the future ocean with high urea emissions and environmentally relevant pH scenarios.

## Introduction

Cyanobacteria are the oldest oxygen-evolving photosynthetic lineage on the earth, and play an important role in global carbon and nitrogen biogeochemical cycles^1,2^. They can use different forms of nitrogen sources, including nitrate, nitrite, ammonium, and some organic nitrogen sources such as urea and amino acids^3-5^. In addition, some cyanobacteria can utilize inert atmospheric nitrogen gas (N_2_) through biological nitrogen fixation, thus providing new nitrogen sources for marine ecosystems ^6,7^. Since phytoplankton including cyanobacteria have different utilization capabilities and preferences for different nitrogen sources, the changes in the form and concentration of nitrogen sources in the ocean may change the composition of marine phytoplankton communities^8-11^. A large amount of excess urea has been emitted into coastal waters by human activities in recent years, resulting in a dramatic increase in urea concentrations in the ocean^12-14^. As a small-molecule organic nitrogen source, urea can be directly utilized by many phytoplankton^15-16^. The increasing input of urea into the ocean could significantly change the community composition of marine phytoplankton, which will have a profound impact on ocean ecosystems and global carbon and nitrogen biogeochemical cycles^17-19^. For instance, it has been reported that dinoflagellates and diatoms benefit from increased urea concentrations in some coastal waters, which might result in occurrence of harmful toxic algal blooms^20-23^. However, so far, a comprehensive understanding of the impact of increasing amounts of urea on the community composition and competitive advantage of marine phytoplankton is still limited.

Acid stress is an environmental factor that cyanobacteria and other phytoplankton need to face. Ecosystem acidification has become a serious problem in the last decades^24^. For instance, human combustion of coal containing sulfur increases the frequency of acid rain by generating sulfur dioxide. Acid rain has been recognized as an important ecological factor with high impact on the coastal ecosystems^25^. Moreover, increasing levels of atmospheric carbon dioxide (CO_2_) due primarily to human usage of fossil fuel cause ocean acidification. The resulting decreased pH values of seawater can alter marine ecosystems^26^. Any reduction of seawater pH will affect the intracellular pH, enzyme activity and energy allocation of related nitrogen metabolism processes of marine phytoplankton, which in turn will affect phytoplankton composition and community succession^27-29^. It has been reported that cyanobacteria can overcome moderate environmental levels of low pH through the expression of particular periplasmic proteins or ion exchange systems, such as Ca^2+^/H^+^ exchanger and Kdp/Ktr uptake systems ^30-32^.

However, little is known about the relationship between urea utilization and acidification adaptation by phytoplankton. It is still unclear how possible interactions between nitrogen utilization and cellular defense mechanisms against low pH may influence the physiology of marine primary producers. In gastrointestinal microorganisms such as *Helicobacter pylori*, the decomposition a large amount of urea produces NH_4_^+^ and OH^-^ and so offers them an important strategy to cope with extremely acidic gastric juices^33^. Here we report that for the first time that cyanobacteria can also use urea to overcome highly stressful low pH conditions, and at the same time very low pH value can alleviate the toxic effect of high concentrations of urea on cyanobacteria. These proof-of-concept laboratory culture experiments under extreme conditions may then point to future experimental strategies to help understand the mechanisms of cyanobacteria responses to eutrophication and acidification interactions under environmental conditions. Urea utilization and acidification adaptation by marine phytoplankton may change the competitive advantages and community composition of marine primary producers, with possible impacts on the marine ecosystem and global biogeochemical cycles.

## Results and Discussion

### Strong interactions between urea tilization and extreme acidification in a coastal cyanobacterium strain

*Synechococcus* sp. strain PCC 7002 (hereafter *Synechococcus* 7002) is a coastal non-nitrogen-fixing unicellular cyanobacterium^34^. It prefers urea as its sole nitrogen source over other nitrogen sources such as NO_3-_ and NH_4_^+^ (Fig. 1A; Fig. S1A-D). When nitrate is used as sole nitrogen source, there is no obvious inhibitory effect on cyanobacteria cells at high concentration (> 12 mM), while high concentrations of ammonium (> 6 mM) or urea (> 12 mM) have toxic effects. Among the three nitrogen sources, *Synechococcus* 7002 showed the highest affinity for urea and obtained the maximum growth rate with moderate concentrations of urea (Fig. 1A; Fig. S1D). We also found that *Synechococcus* 7002 can remain dormant for more than three months during long-term nitrogen deficiency, and once new nitrogen sources are available, they can resume growth (data not shown). Our data indicated that the addition of urea can more quickly restore the growth of nitrogen-deficient *Synechococcus* 7002 than the addition of ammonium or nitrate (Fig. S1E and S1F). Under laboratory culture conditions (see the Materials and Methods), *Synechococcus* 7002 required 12 mM nitrate to reach maximum growth rates (Fig 1A, 1B, and Fig. S1A). Further increases of nitrate concentration did not increase nor inhibit the growth rate of *Synechococcus* 7002. However, even at the saturated nitrate concentration, the addition of urea (6 mM, 12 mM nitrogen source, since one urea molecule has two nitrogens) can further increase the growth rates of *Synechococcus* 7002. In contrast, the addition of ammonium at the same concentration of nitrogen (12 mM) inhibited its growth (Fig. 1B and Fig. S2A). These results indicated that the preference of *Synechococcus* 7002 for urea did not depend on the concentration of nitrate, which is different from the utilization of ammonium. It has been previously shown that high concentration of nitrate generally reduces the dependence of phytoplankton on ammonium^35^.

**Figure 1.**
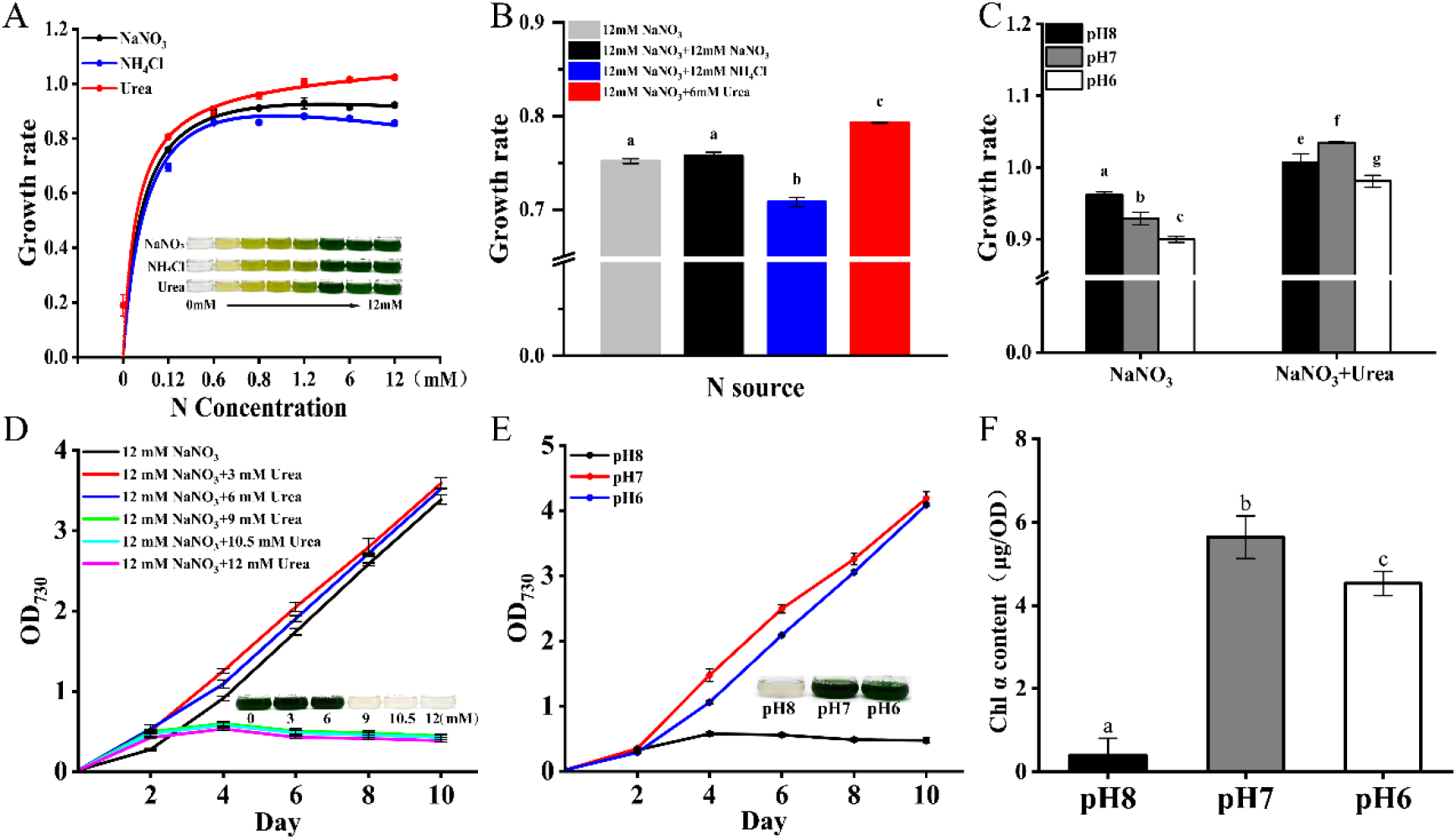
(A) Fitted curves of growth rates of *Synechococcus* 7002 with three nitrogen sources (NaNO_3_, NH_4_Cl and urea) at a range of concentrations (0 to 12 mM nitrogen). Shown is the fourth day of the culture. (B) Growth rates of *Synechococcus* 7002 with saturating NaNO_3_, with and without three additional nitrogen sources. (C) Growth rates of *Synechococcus* 7002 at three different pH conditions, with or without urea (12 mM NaNO_3_, and 12 mM NaNO_3_ plus 6 mM urea). (D) Growth curves of *Synechococcus* 7002 under saturated N concentration (12 mM NaNO_3_) and different concentrations of urea. (E and F) Growth curves and Chl *a* content of *Synechococcus* 7002 at different pH conditions with a high nitrogen concentration of urea (24 mM). The error bar in all the experiments represents the standard deviation between the three replicates, and the significance analysis is marked with lowercase letters on each bar, with different letters representing significant differences (*P* < 0.05).

When pH values of the culture conditions decreased from a seawater-relevant value 8.0 to an extreme acidic level of 6.0, the growth of *Synechococcus* 7002 was proggressively inhibited as expected (Fig. 1C and Fig. S2B). However, the addition of urea alleviated the inhibitory effect of this highly stressful acidification treatment. As shown in Fig. 1C, when urea was added, the decreasing trend of growth rate with the decrease of pH value changed. Although the growth rate at extremely low pH (6.0) is still the lowest among the three pH conditions, the negative effect of acidification at pH 7.0 disappeared (Fig. 1C and Fig. S2B). The growth rate at pH 7.0 is even higher than that at standard seawater pH levels of 8.0 when urea was added, suggested that urea is playing a positive role in counteracting acidification. This phenomenon was also found in several eukaryotic phytoplankton, such as the diatom *Thalassiosira pseudonana* and the dinoflagellate *Karenia mikimotoi* (Fig. S3). The distribution of this phenomenon among different marine phytoplankton is still under investigation. As mentioned above, very high concentration of urea (higher than 12 mM urea) caused a toxic effect on *Synechococcus* 7002, including an obvious bleaching phenotype and photosynthetic damage (Fig. 1D and Fig. S4). This inhibition phenotype under high concentrations of urea at standard pH conditions is consistent with previous reports^36^. However, lowering culture pH value from 8.0 to 6.0 can obviously alleviate the toxic effects of high concentrations of urea (Fig. 1E and Fig. S4). As shown in Fig. 1E and 1F, *Synechococcus* 7002 cannot grow at all with a very high concentration of urea (24 mM) at pH 8.0, showing a completely bleaching phenotype. However, it can grow well at much lower pH values (7.0 and 6.0) with the same concentration of urea. This result further suggests there is a strong interaction between urea utilization and acidification adaptation in cyanobacteria.

### Urease allows cyanobacteria to cope with extreme acidification

Urea utilization by cyanobacteria depends on an important intracellular enzyme, urease. Urease is widely found in prokaryotes, fungi, and archaea. It requires a nickel cofactor to catalyze the breakdown of urea into NH_4_^+^, CO_2_ and OH^- 37^. In cyanobacteria, urease is generally composed of seven different subunits UreA ∼ UreG, and the loss of any subunit results in inactivation of the ability to break down urea (Fig. S5 and Fig. S6). We constructed mutants of urease subunits and found that they can grow normally with NaNO_3_ or NH_4_Cl as nitrogen sources, but cannot grow on urea as sole nitrogen source as expected (Fig. 2A-C; Fig. S6). Interestingly, all these urease mutants showed more sensitivity to extreme acidification conditions than the wild type, with mixed nitrogen sources (NaNO_3_ plus urea). Taking *ureC* and *ureG* mutants for examples, the mutants lost the ability to overcome acidification and showed a sensitivity to low pH conditions (Fig. 2D). Unlike the wild type showing the highest growth rate under pH 7.0, the growth rate of the *ureC* and *ureG* mutants decreased as pH was lowered from 8.0 to 6.0 (Fig. 2D), which is similar to the phenotype of the wild type cultured without urea. These results confirmed that urea utilization and urease are crucial to adaptation to very high levels of acidification in *Synechococcus* 7002. We further investigated the urease activity of the wild-type strain of *Synechococcus* 7002, and found that under pH 7.0 and pH 6.0 conditions the urease activities are significantly higher than that under the pH 8.0 condition (Fig. 2E). This suggests that a high urease activity is required by *Synechococcus* 7002 to adapt to highly acidified conditions.

**Figure 2.**
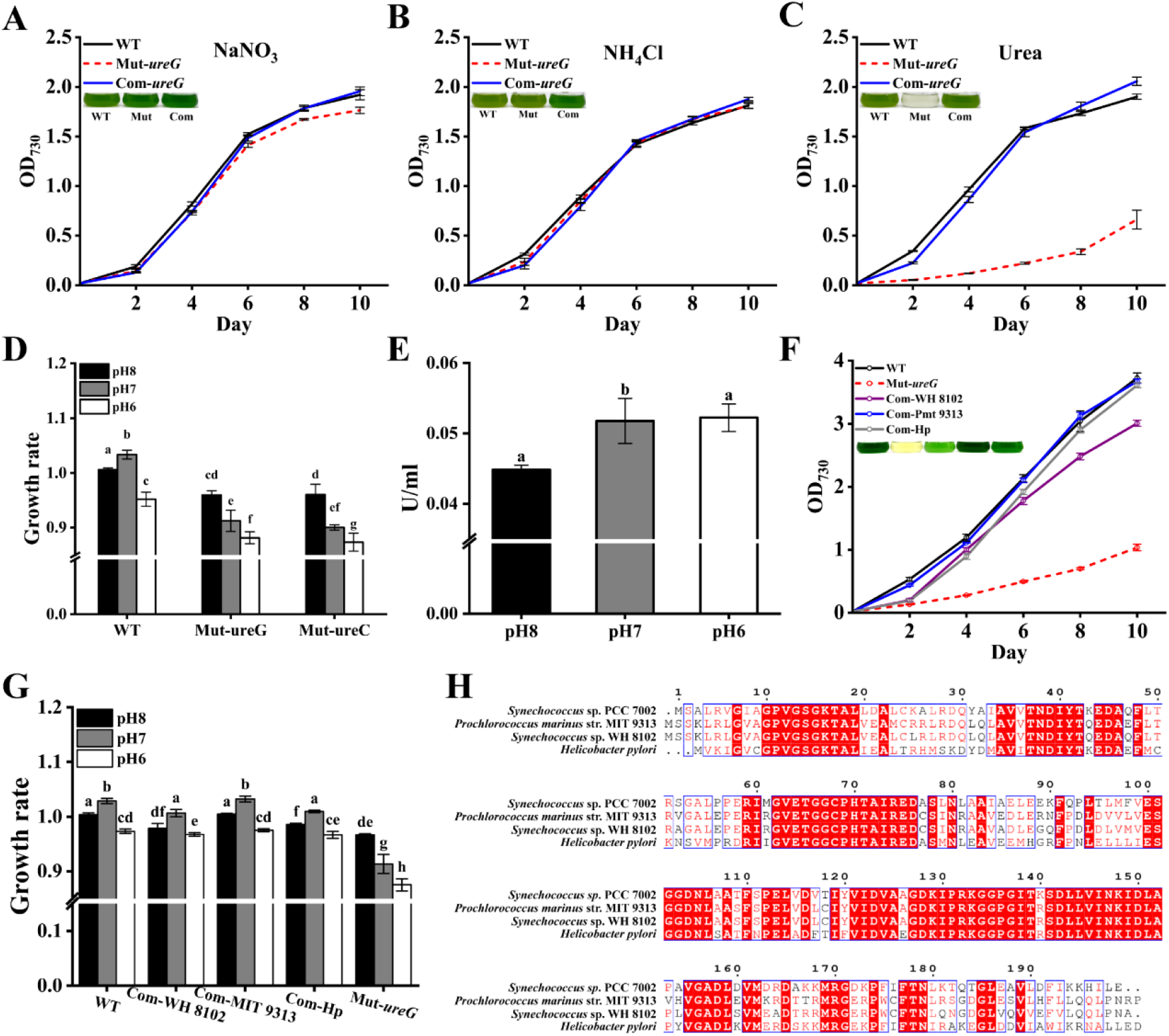
(A, B and C) The growth curves of the wild-type (WT) and *ureG* mutant strain (Mut) and complemented strain (Com) of *Synechococcus* 7002 under 12 mM nitrogen concentrations from different nitrogen source (NaNO_3_, NH_4_Cl and Urea). The inserted photos show the strains on the 4th day of incubation under the corresponding conditions. (D) Growth rates of *Synechococcus* 7002 wild-type and mutant strains (*ureC* and *ureG*) at different pH with mixed nitrogen sources (12 mM NaNO_3_ plus 6 mM Urea). (E) Urease activity of *Synechococcus* 7002 under mixed nitrogen source (12 mM NaNO_3_ + 6 mM Urea) and different pH conditions. (F) Growth curves of *Synechococcus* wild-type (WT), *ureG* mutant strain, and complementary strains under pH 8.0 conditions. In complementary strains, homologs of *ureG* from *Helicobacter pylori* (Com-Hp) and *Synechococcus* sp. WH 8102 (Com-WH 8102) and *Prochlorococcus* sp. MIT 9313 (Com-Pmt 9313) was heterologously expressed in the *ureG* mutant of *Synechococcus* 7002. (G) Growth rates of the *Synechococcus* 7002 wild type and complemented strains under different pH conditions and mixed nitrogen sources (12 mM NaNO_3_ + 6 mM Urea). Com-WH 8102, Com-MIT 9313, and Com-Hp represent the complemented strain of *Synechococcus* 7002 by heterologous expressing UreG homologs of *Synechococcus* sp. WH 8102, *Prochlorococcus marinus* str. MIT 9313 and *Helicobacter pylori*, respectively in the *Synechococcus* 7002 Mut-*ureG* mutant. (H) The amino acid sequence alignment of UreG from *Synechococcus* 7002, *Synechococcus* sp. WH 8102, *Prochlorococcus marinus* str. MIT 9313 and *Helicobacter pylori*. The error bar represents the standard deviation between the three replicates, and the significance analysis is marked with lowercase letters on each bar, with different letters representing significant differences (*p*<0.05).

We confirmed that these phenotypes of the mutants were indeed caused by the inactivation of urease encoding genes rather than a second point mutation, since the complement of the target genes back into the mutants restores them to phenotypes comparable to the wild type (Fig. 2A-C, Fig. S7, taking Mut-*ureG* and Mut-*ureC* for examples). It has been reported that some acidophilic heterotrophic bacteria such as *Helicobacter pylori* and *Klebsiella pneumoniae* can raise intracellular pH by degrading urea and producing a large amount of NH_3_ (NH_4_^+^ and OH^-^) to overcome extremely low pH conditions^38^. So far, this strategy of fighting acidification has mainly been reported in a few oral and gastrointestinal microbes^38^. Using transgenic methods and bioinformatic analyses, we found that the ureases from *Helicobacter pylori* and different marine cyanobacteria species play a similar function. The heterologous expression of UreG homologs from *Helicobacter pylori* and some open-ocean cyanobacterial strains (*Synechococcus* sp. WH 8102 and *Prochlorococcus* sp. MIT 9313) can also restore the phenotypes of the Mut-*ureG* strain of *Synechococcus* 7002, including both the capability to utilize urea and adaptation capability to low pH conditions (Fig. 2F and 2G). These results implied that the roles of ureases from different microorganisms are conserved (see Fig. 2H for the amino acid sequence alignment and Fig. S8 for phylogenetic tree of UreC and UreG). The ability of urea to compensate for acidification is likely to exist in much wider range of species than previously considered, far beyond gastrointestinal microbiota.

### The distribution of cyanobacteria containing urease is correlated with high urea emissions in coastal waters

Because urea concentration was not routinely measured in most previous research voyage studies, specific data for urea concentrations in many global ocean areas are still lacking. However, in the past 20 years the annual amount of urea used globally has increased about four times, from 40 million tons to 180 million tons (Fig. S9A). Our results show that cyanobacteria with urease genes may be favored in coastal waters with high urea emissions. By comparing the metagenomic data from TARA ocean sites (which lacks data from Eastern Asia ocean regions), we found that cyanobacteria with urease genes are distributed in coastal waters close to the regions with high urea emissions, especially in the marginal seas of the Indian Ocean and the Red Sea (Fig. 3). According to the latest data published by Food and Agriculture Organization of the United Nations(FAO) (https://www.fao.org/faostat/en/#data/RFB/visualize) and previous reports (Fig. S9B and Table S2), all the countries near these coastal waters are reported to use large amounts of urea for agricultural production (Table S2). Using different subunits of cyanobacteria urease or the urease of eukaryotic phytoplankton (such as dinoflagellates) for this analysis, the resulting distribution of cyanobacteria with urease (data not shown) are similar to that obtained using the urease genes of *Synechococcus* 7002. All the sites with abundant distributions of cyanobacterial urease are coastal waters, except the sites located in the southeastern Pacific Ocean (Fig. 3). We found that these sites are actually near South Eastern Pacific islands, where significant anthropogenic discharges of urea with agricultural activities and large amount of groundwater nitrogen inputs have been reported^39-42^.

**Figure 3.**
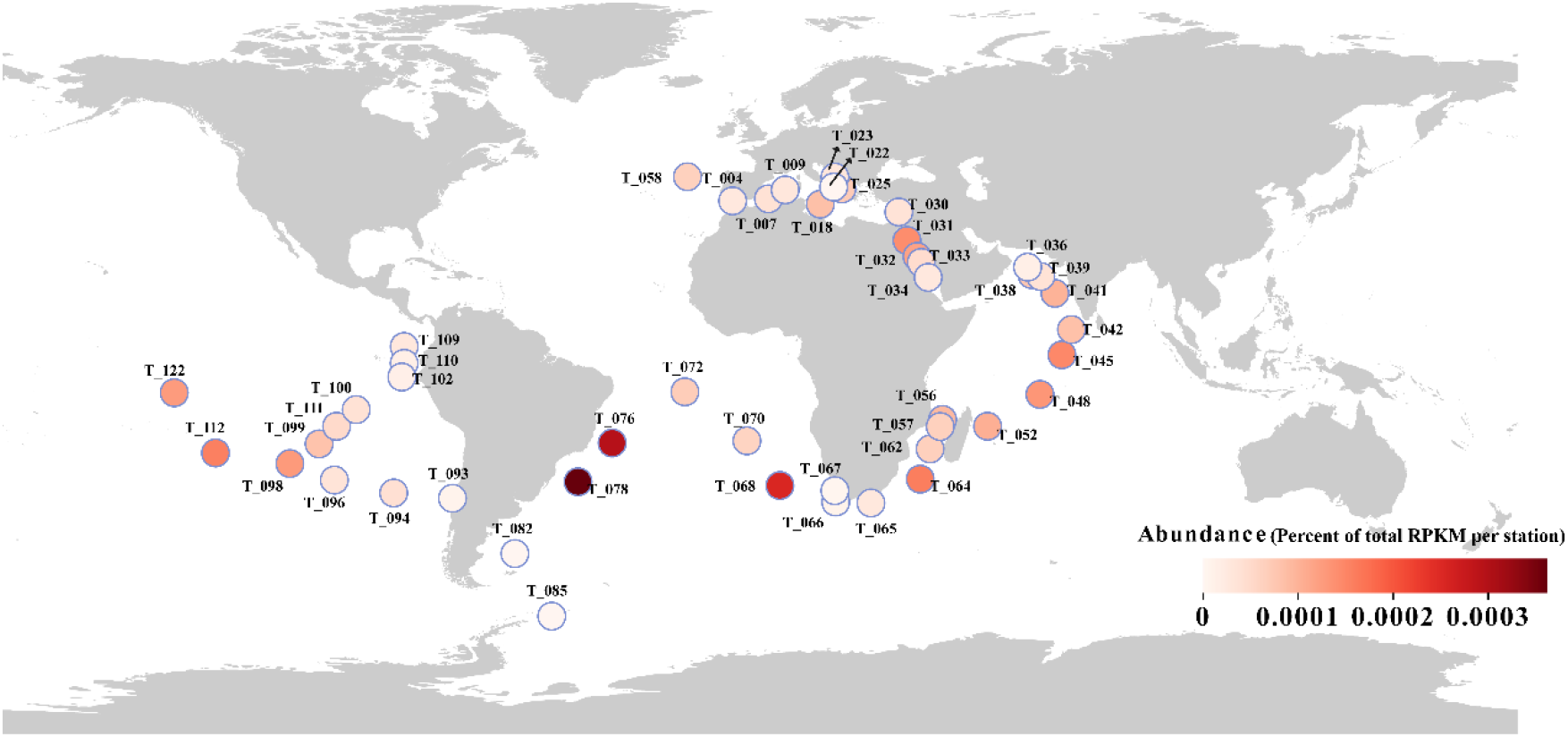
(A) Global distribution of cyanobacteria with urease in the surface water and DCM layers of the Tara Ocean sites analyzed with the large subunit of urea UreC amino acid sequence. T_XXX (e.g., T_078) in the figure represents the station number, and the color shades of the circles represent the abundance UreC of cyanobacteria in the specific site. The composition and percentage of cyanobacteria are shown in the supplementary data Fig. S10.

The large influx of undegraded urea into coastal waters is probably one of the reasons for the abundance of cyanobacteria containing urease genes. Among the cyanobacterial species with urease genes found in TARA Ocean sites, the proportion of *Prochlorococcus* spp. is the highest (more than 70%), followed by *Synechococcus* (Fig. S10). The distribution characteristics of these cyanobacteria with urease in the coastal waters with high concentration of urea is likely related to their better urea utilization and metabolism capacity. Some marine cyanobacteria such as *Synechococcus* sp. WH 7803 have been reported to lack urease and the ability to use urea^43^. According to our physiological data, these cyanobacteria or phytoplankton without urease are likely to lose their competition advantages in these environments. At the same time, different phytoplankton exhibited different urea utilization capability and preference, the competitive advantages and composition of marine phytoplankton is likely to be changed dramatically due to the urea input into the ocean. Our experiments at extreme pH levels suggest that these mitigation mechanisms mediated by urea utilization should be investigated in further experiments at more moderate pH levels that are relevant to near-future anthropogenic ocean acidification by atmospheric CO_2_ and acid rain in coastal waters (e.g., pH values of 7.7-7.9). It is possible that even at these much lower levels of acidity, the coupling effect of increased acidification and urea concentration may provide urea-utilizing phytoplankton populations with an advantage in the future ocean.

