## Supplementary material for "Cyanobacteria using urea as a nitrogen source can overcome acid stress": Supplemental file.docx

**Materials and Methods**

**Strains, culture conditions and general methods**

*Synechococcus* sp. strain PCC 7002 (hereafter *Synechococcus* 7002) wild type, mutant strains and complemented stranis were cultured in A^+^ medium under continuous light 100 μmol photons m^-2^ s^-1^, at 30 °C, with a rotatory shaker at 110 rpm. All growth media, buffers and solutions used in the experiments were autoclaved or filter sterilized. According to the requirements of the mutants or complemented strains, kanamycin (25 mg·ml^-1^ ) or spectinomycin (20 mg·ml^-1^) was added into the medium. Standard A^+^ medium was adjusted to pH 8.0 by Tris-HCl buffer, and low pH medium was adjusted to pH 7.0 and pH 6.0 buffered with 1 M HEPES Buffer and 0.5 M MES Buffer, respectively, for detail refer by our prevous study of Jiang et al. 2015 ^1^. The growth of all cyanobacterial strains was assessed by measuring turbidity (OD_730_) from three independent biological replicates every two days. All strains were cultured at the same initial OD_730_ value of 0.02 with three replicates. Growth rates were calculated according to formula: LN[OD_730(Dayt2)_ /OD_730(Dayt1)_]/(Day_t2_-Day_t1_) based the linear logarithmic phase. To determine chlorophyll *a* (Chl *a*) content, the absorption peaks at 648.6 and 664.1 nm of 95% ethanol extracts were measured using an ultraviolet-visible spectrophotometer TU1810 (PERSEE, China) and calculated according to the formula: Chl *a* =13.36×A_664.1_−5.19×A_648.6_, for details of the method refer by Liu *et al* (2022)^2^. The culture conditions for eukaryotic algae (*Thalassiosira pseudonana* and *Karenia mikimotoi*) were continuous light 30 μmol photons m^-2^ s^-1^, at 22 °C, in f/2 medium. The growth of eukaryotic algae was determined by cell counting using a CytoFLEX flow cytometer (Beckman Coulter, America). See Table S3 for the strains used in this experiment. As for the measurement of urease activity of *Synechococcus* 7002, the amount of NH_3_-N produced by urease hydrolysis of urea was used by indophenol blue colorimetric method with the cells in exponential phase. The cells at the 4^th^ day of growth were collected and diluted, and the urease activity of the 1 ml solutions was measured by the urease activity detection kit (Solarbio, BC4115). All data are given as mean and standard deviation of three biological replicates, statistics analysis method was used with an independent sample T-test. We have marked Significant differences with different letters (*p* < 0.05).

**Construction of mutant and complemented strains**

Taking Mut-*ureG* strain as an example, using the DNA of wild-type *Synechococcus* 7002 as a template, PCR amplification was carried out with *ureG*-up-F/*ureG-*up-R and *ureG*-DN-F/*ureG*-DN-R primers respectively to obtain the upstream homologous segment of *ureG* gene, *ureG*-UP, and the downstream homologous segment, *ureG*-DN. The recombinant plasmid NBU-KAN-MUT-*ureG* was obtained by inserting the *ureG*-UP fragment and *ureG*-DN fragment into the upstream and downstream of the kanamycin resistance gene (Kan^R^) of a pUC-MCS-KAN-MCS plasmid vector, respectively and keep the gene sequences in the same orientation (Fig. S5 B). Then the recombinant plasmid NBU-KAN-MUT-*ureG* was transformed into the wild-type *Synechococcus* 7002 to obtain the mutant of UreG (Mut-*ureG*). For complemented strains, the exogenous gene from different species was heterologous expressed under a promoter of porin gene (A2813) of *Synechococcus* 7002. As shown in Fig. S5C, the expression cassete of the porin promoter, an exogenous gene, and an spectinomycin resistance gene (Sp^R^) fragment (Fig. S5E) was inserted into the neutral site of *Synechococcus* 7002 genome (A0935-0936 location site) ^3^. The transformation method refers to the description of Williams et al (1988) and our previous studies ^4^. The primers used in this study are shown in Table S1.

**Measurement of chlorophyll fluorescence parameters**

Photosynthetic parameters were measured using a FluorPen FP 100 fluorometer (Photon System Instrument, Czech Republic). Maximum photochemical efficiency of photosystem II (F_v_/F_m_) was measured with the cells at logarithmic growth stage by calculating the minimum fluorescence value F_0_ (minimal fluorescence emission, open PSII centres) after 20 min of dark acclimation and F_m_ (maximal fluorescence emission, closed PSII centres) after adding DCMU (dichlorophenyl dimethylurea) and the caltulation fomula is F_v_/F_m_=(F_m_-F_0_)/F_m_. The maximum relative electron transport rates (rETR_max_) and the light-saturation parameter (I_k_) were also measured with cells at logarithmic growth stage and resulting data were fitted using origin pro 2021 software acording the methods of Ralph *et al* (2005) ^5^.

**Bioinformatics analysis**

As for the distribution of cyanobacteria urease genes in the ocean, the UreC sequecence from *Synechococcus* 7002 and other cyanobacteria was alignment on the Tara Ocean website (<http://tara-oceans.mio.osupytheas.fr/ocean-gene-atlas/>). The abundance matrix data from deep chlorophyll maximum (DCM) layer and surface sample fractions of cyanobacterial at each site were selected from the alignment results for analysis and mapping and the detailed operation process was performed as described by Vernette *et al* (2022)^6,7^. Sequence alignment using the software MEGA-X, results were visualized on the image processing website ESPrirt 3.0 ^8^. Phylogenetic trees were conducted based on the multiple sequence alignments using the Neighbor-Joining method in MEGA-X with 1000 bootstrap replicates. Phylogenetic trees and map were visualized on the image-processing website Chiplot (<https://www.chiplot.online/chitree.html>).

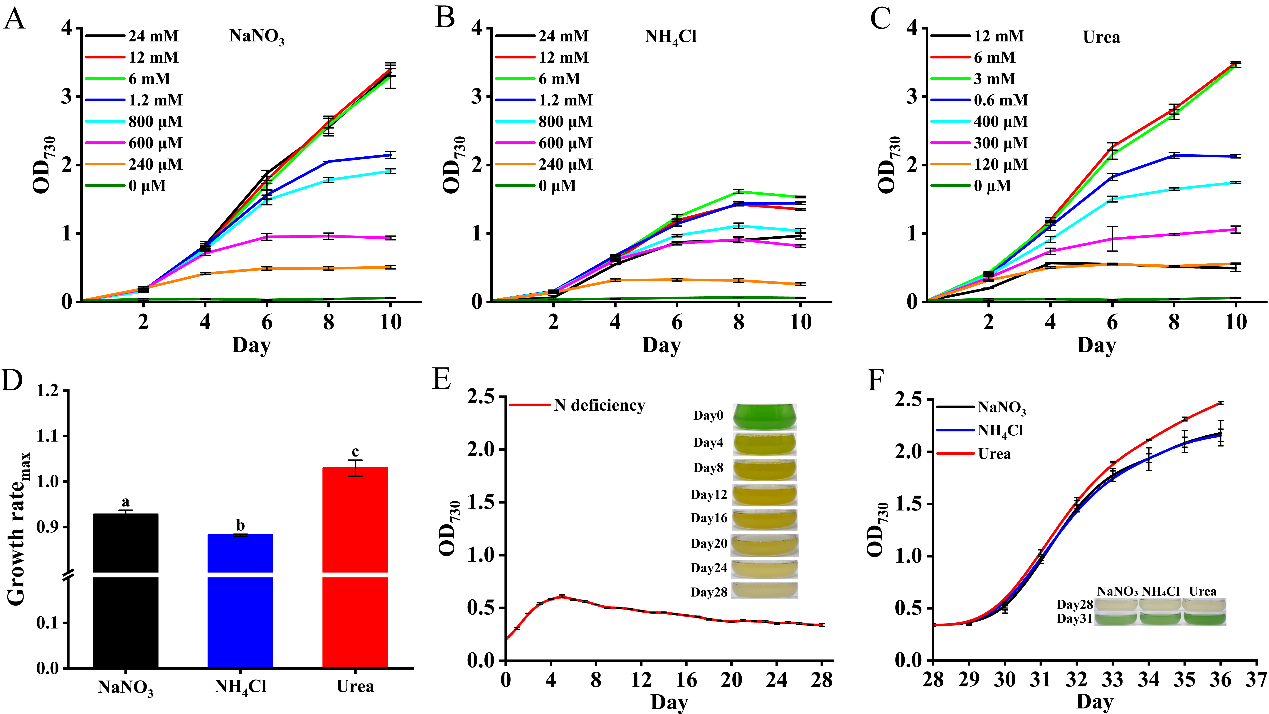

Figure S1 The physiological characteristics of *Synechococcus* 7002 under different nitrogen conditions. A, B ,C, are the growth curves of *Synechococcus* 7002 under different concentration of NaNO_3_, NH_4_Cl and Urea as nitrogen sources (0 to 24 mM nitrogen concentrations). D shows the maximum growth rate of *Synechococcus* 7002 with different nitrogen sources. E shows the growth curve of *Synechococcus* 7002 after long term of nitrogen deficiency incubation. F is the growth curve of *Synechococcus* 7002 when the long term of nitrogen deficienct culture (28 days) was added with 1.2 mM of different nitrogen sources (NaNO_3_, NH_4_Cl and Urea). The error bar represents the standard deviation between the three replicates, and the significance analysis is marked with lowercase letters on each bar, with different letters representing significant differences, (*p*<0.05).

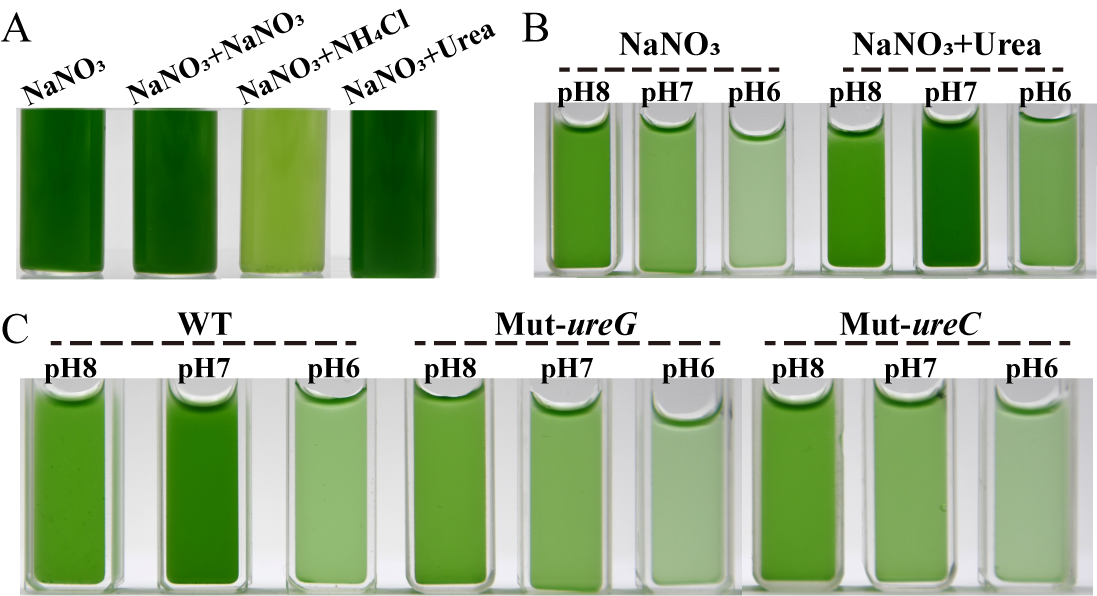

Figure S2 Photographs of *Synechococcus* 7002 strains under different culture conditions. A，B, and C corresponds to the growth photos of strains on the fourth day shown in Figure 1B, 1C, and 1D in the main Text, respectively.

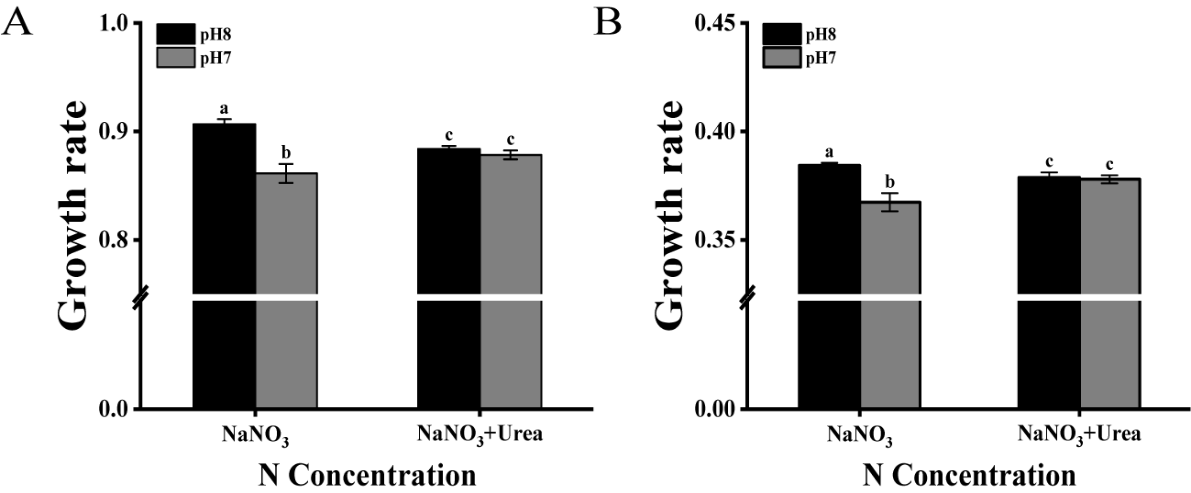

Figure S3 The growth rates of *Thalassiosira pseudonana* (A) and *Karenia mikimotoi* (B) in f/2 medium with different nitrogen sources (880 μmol NaNO_3_ and 880 μmol NaNO_3_ plus 440 μmol Urea) and different pH conditions (pH 8.0 and 7.0) . The error bar represents the standard deviation between the three replicates, and the significance analysis is marked with lowercase letters on each bar, with different letters representing significant differences, (*p*<0.05).

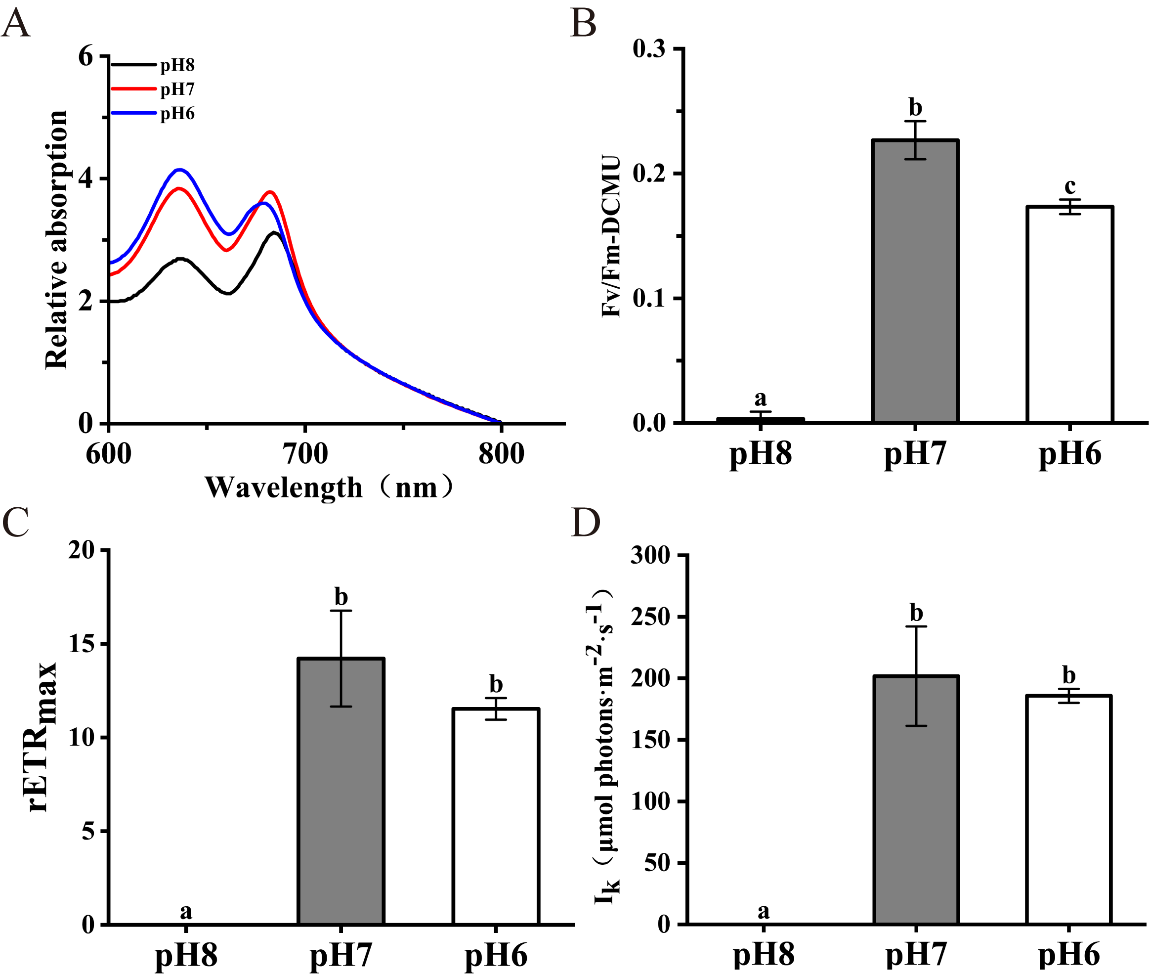

Figure S4 Whole-cell absorption spectrum (A), photosynthetic parameters (B, C, D) of *Synechococcus* 7002 at high concentration of urea (12 mM NaNO_3_ plus 12 mM Urea) under different pH conditions (8.0, 7.0 and 6.0). The error bar represents the standard deviation between the three replicates, and the significance analysis is marked with lowercase letters on each bar, with different letters representing significant differences, (*p*<0.05).

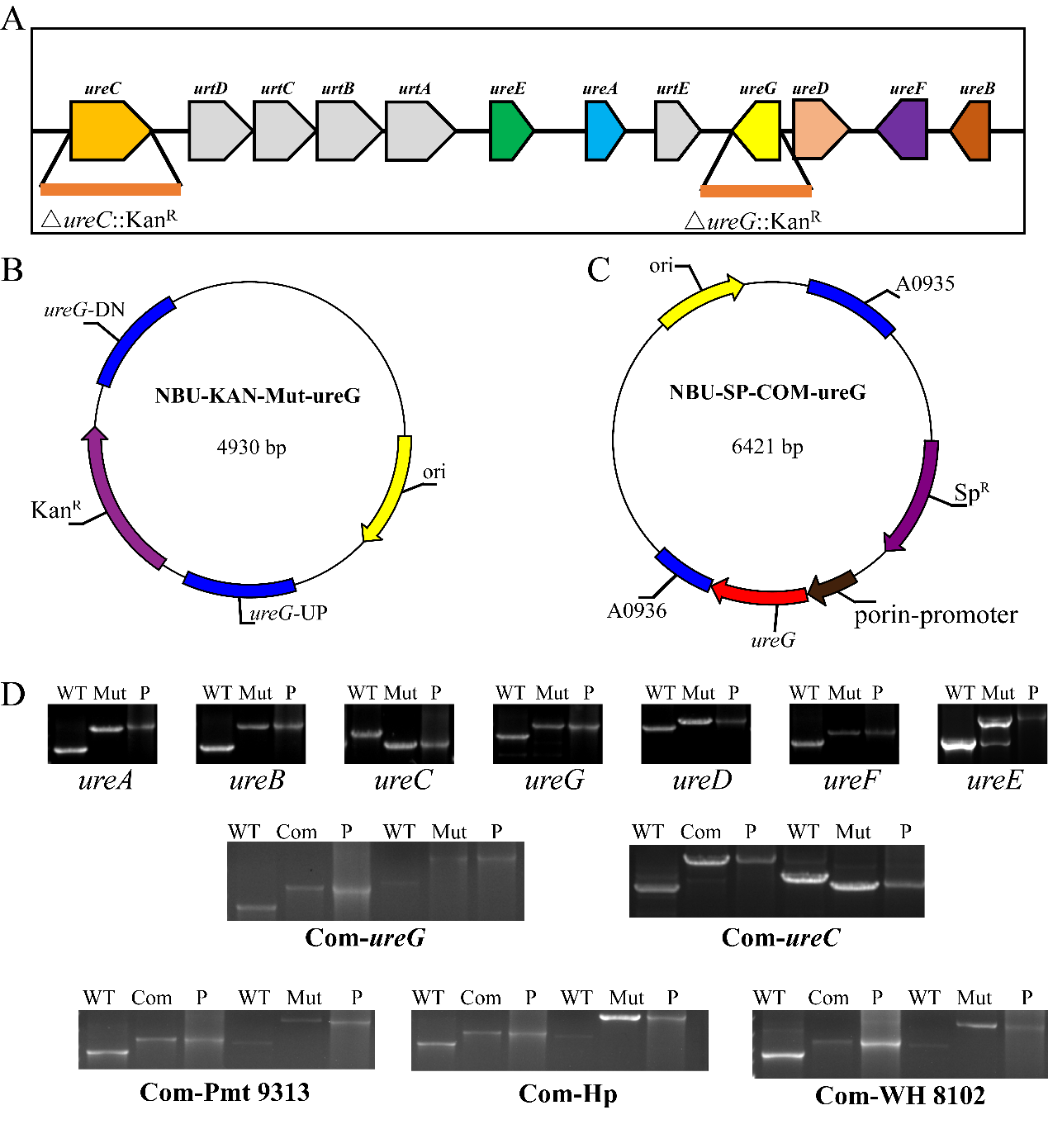

Figure S5 The subunit composition and genome location of urease coding gene, the construction of knockout and complemented plasmids, and PCR detection of the strains of *Synechococcus* 7002. A shows the location of the encoding genes of different urease subunits in the chrosome of *Synechococcus* 7002, as well as the knockout pattern (taking with *ureC* and *ureG* as examples). B and C, schematic diagram of construction of plasmid for knockout and complementary strains (taking Mut-*ureG* as an example). D shows the PCR detection confirming the succesful construction of the coresponding mutants and complemented strains of *Synechococcus* 7002. PCR primers used in this study are shown in Table S1. In each photo, from left to right is the results of PCR with the genome of *Synechococcus* 7002 (WT) the mutants (Mut), and the knockout plasmids (P) with template, respectively. The *ureE* gene was not completely knocked out.

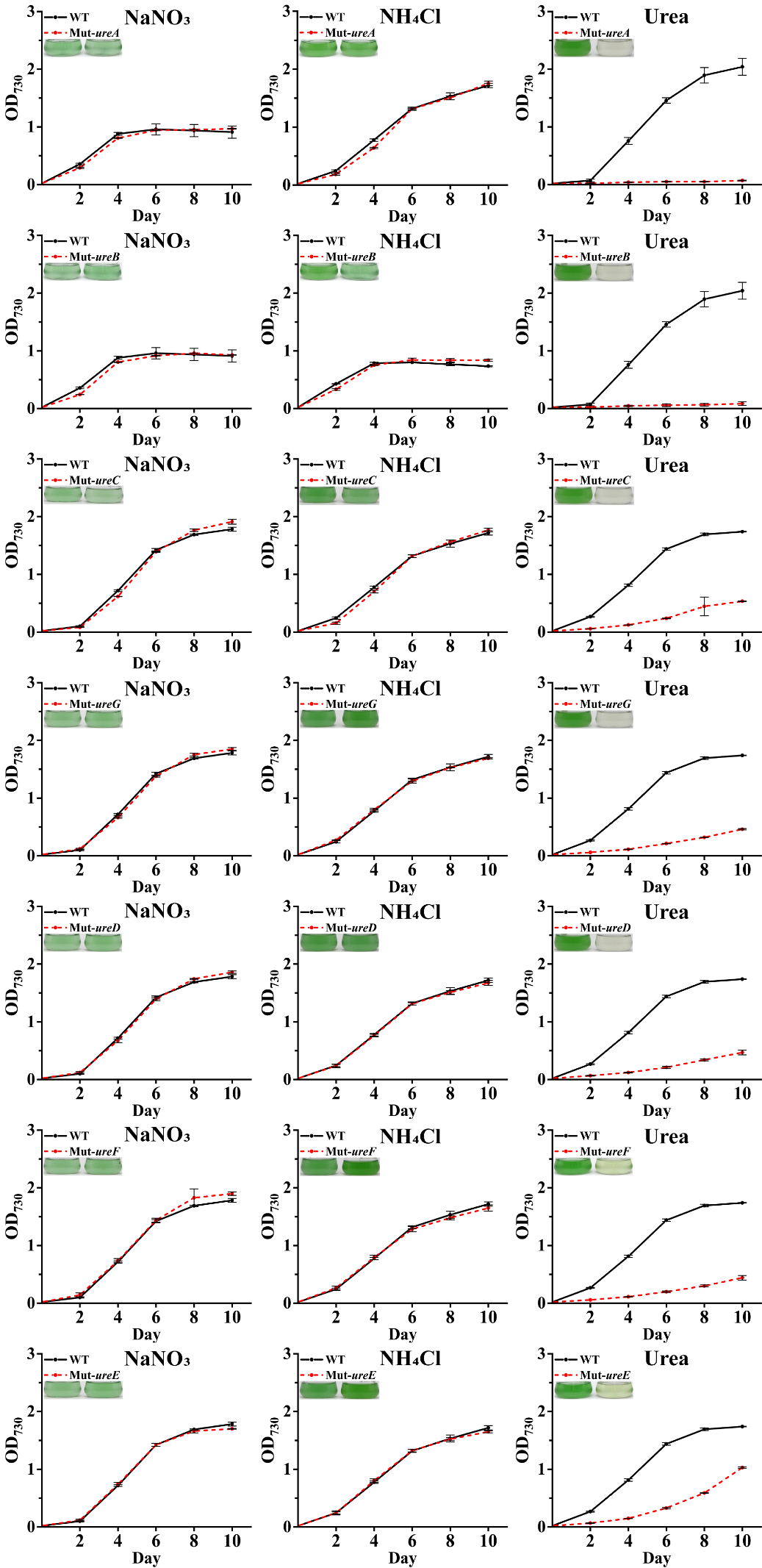

Figure S6 The growth curves of seven different mutants of *Synechococcus* 7002 urease subunits (Mut-*ureA*, Mut-*ureB*, Mut-*ureC*, Mut-*ureG*, Mut-*ureD*, Mut-*ureF*, Mut-*ureE*) with three sole nitrogen sources (1.2 mM NaNO_3_, 1.2 mM NH_4_Cl and 0.6 mM Urea). All strains were incubated for 4 days without nitrogen source before the experiment. The inserted photos show the strains on the 4^th^ day of culture under the corresponding conditions. The error bar represents the standard deviation between the three replicates.

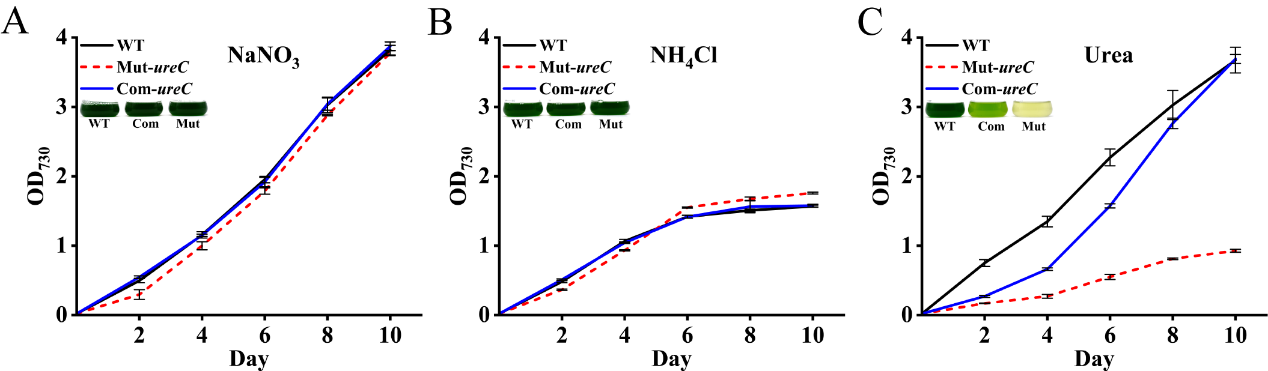

Figure S7 The growth curves of WT and *ureC* mutant strain and complemented strain of *Synechococcus* 7002 under 12 mM nitrogen concentration of different nitrogen source (NaNO_3_, NH_4_Cl and Urea) The inserted photos show the strains on the 4th day of incubation under the corresponding conditions. The error bar represents the standard deviation between the three replicates.

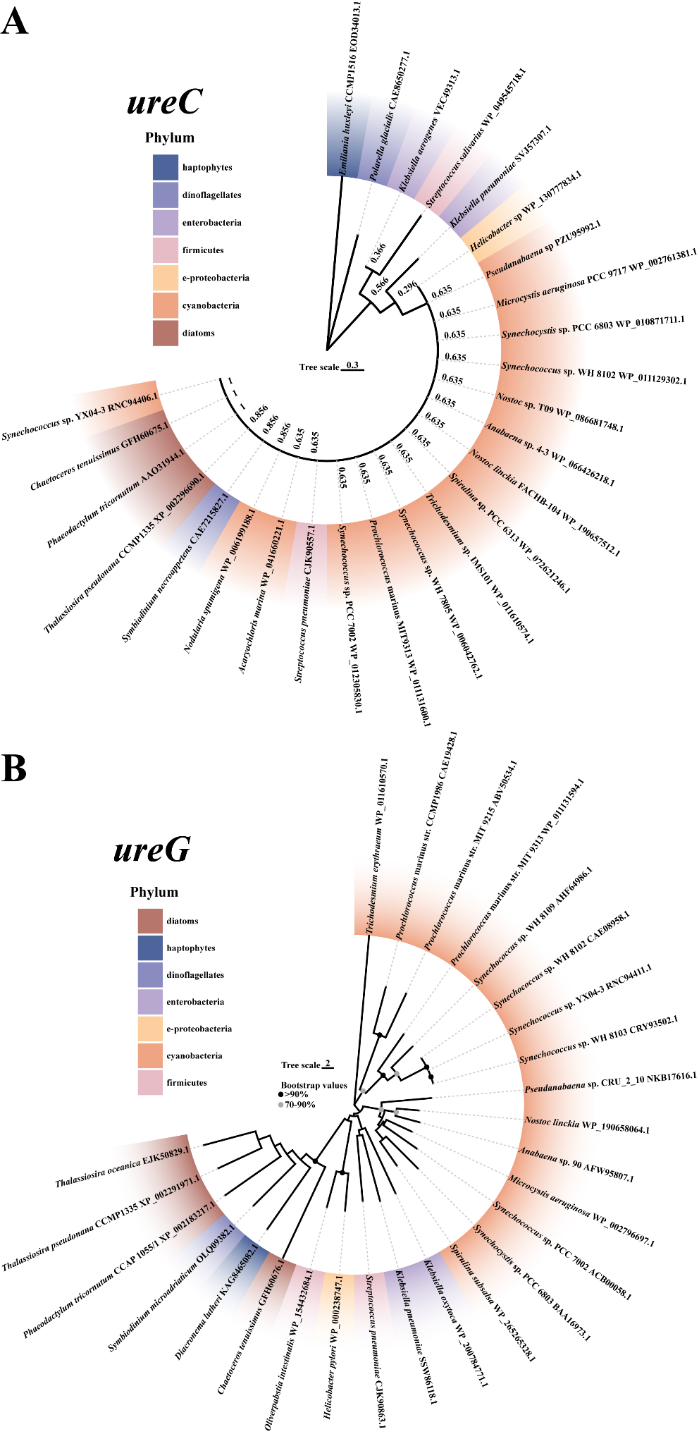

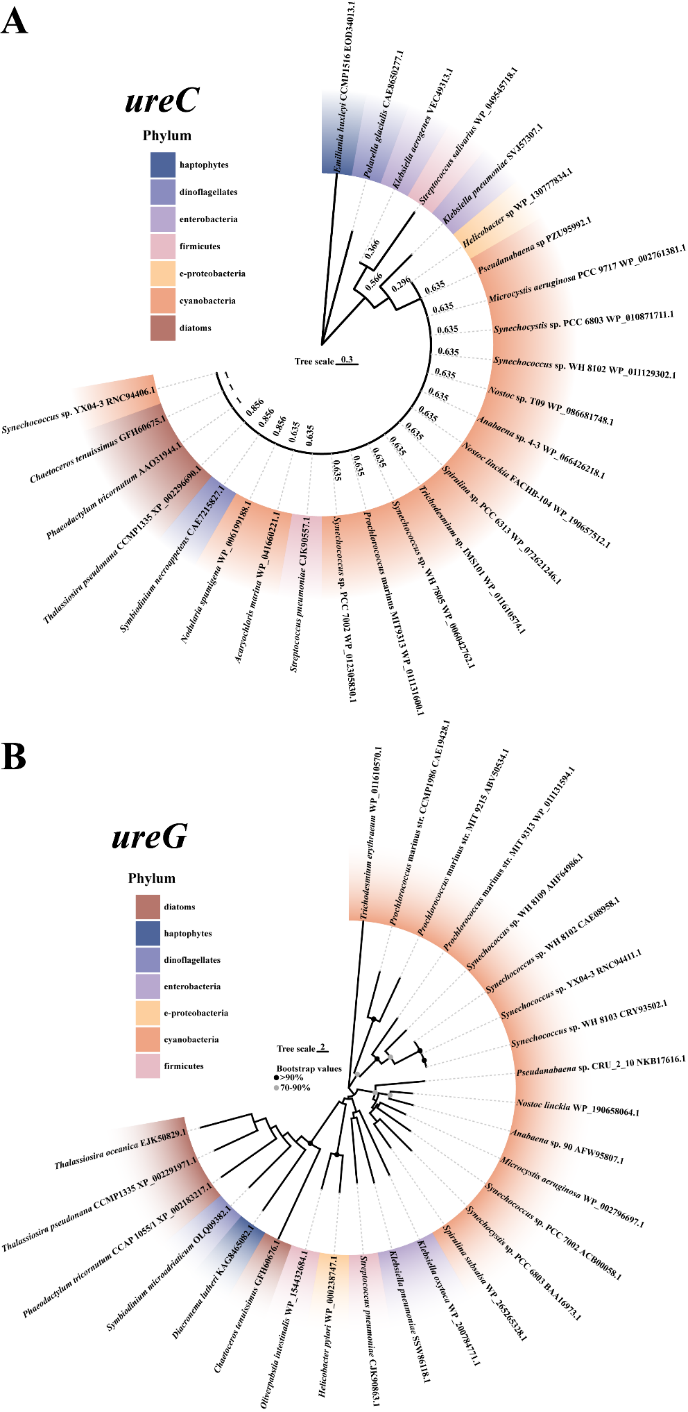

Figure S8 Unrooted phylogenetic tree of UreC and UreG constructed from species of different phylum such as pathogenic bacteria, cyanobacteria, diatoms, and dinoflagellate et al. Using Neighbour-joining method. The species of different phyla in the figure are distinguished by different background colors.

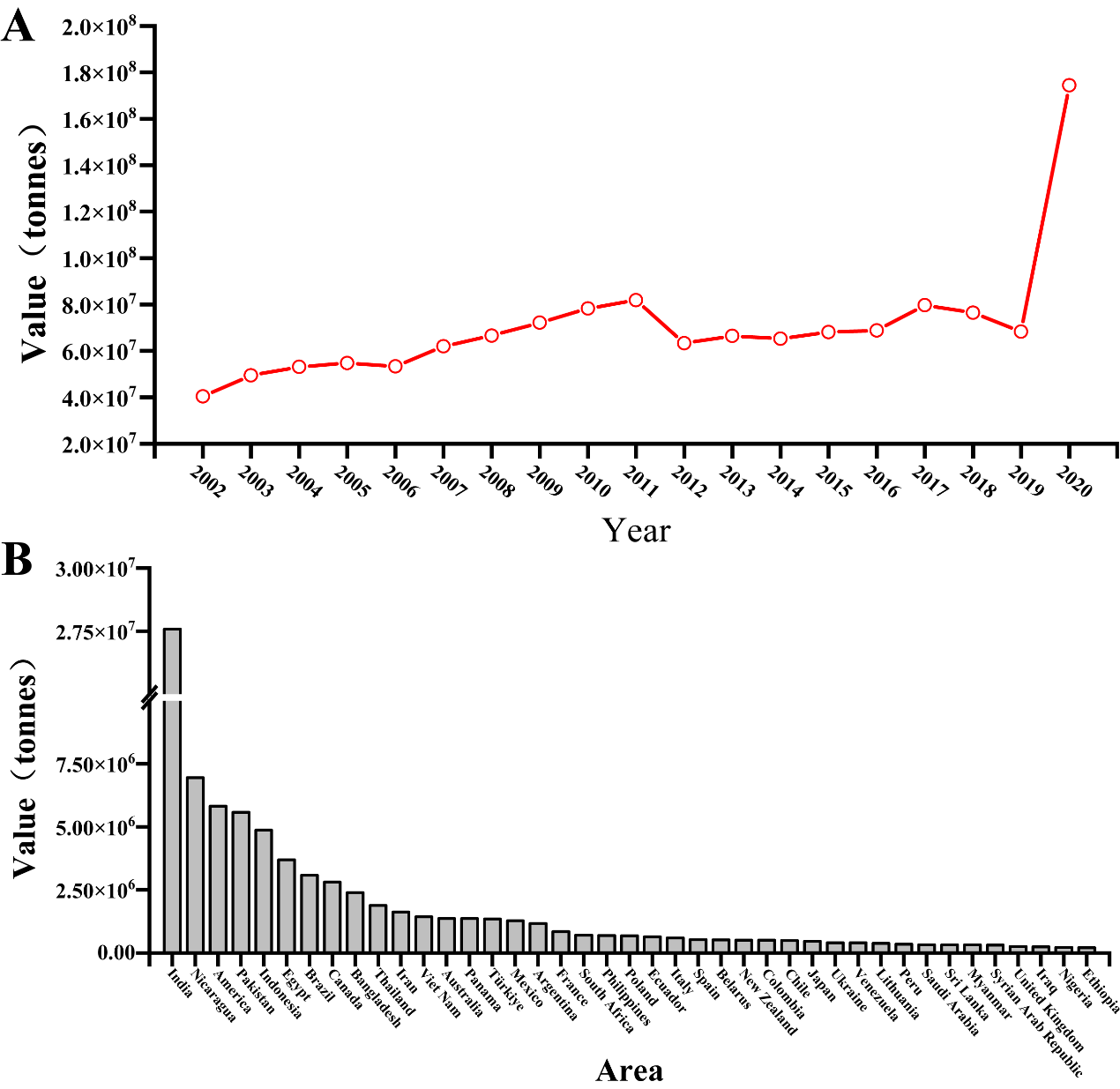

Figure S9 Global annual use amounts (A) and annual use amounts of urea in some countries or regieons (B) reported by FAO.

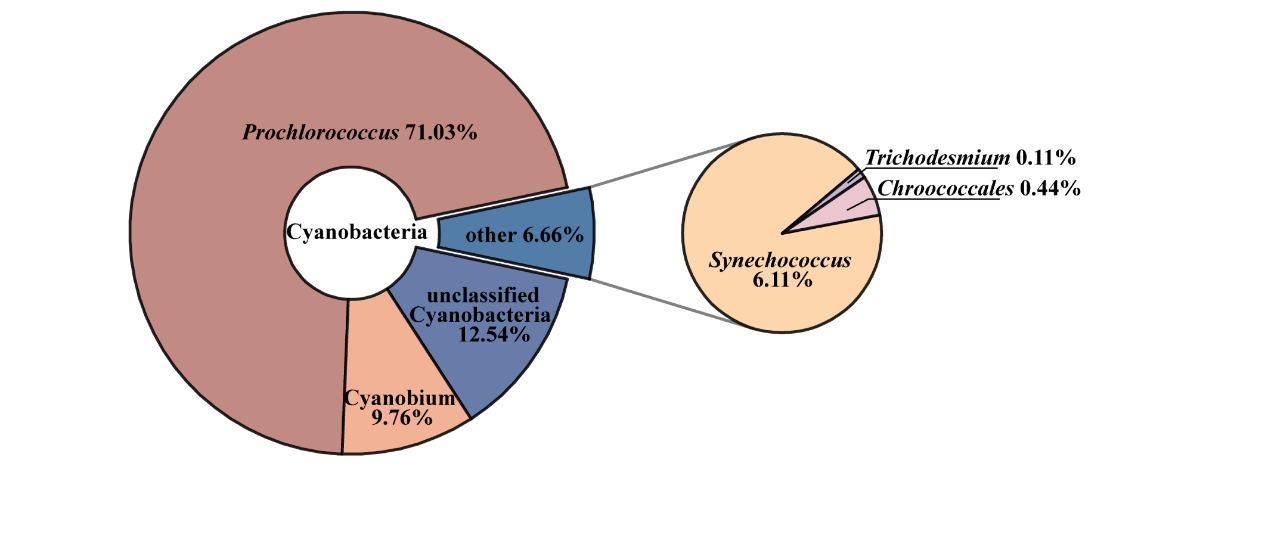

Figure S10 All cyanobacterial species and proportions detected in Figure 3, with the largest proportion of *Prochlorococcus* (>71%), followed by unclassified cyanobacteria and some other cyanobacteria, including *Synechococcus* sp. WH 7805 and *Synechococcus* sp. WH 8016 and other unclassified *Synechococcus*.

Table S1 Primers used in this study.

| Primer name | Sequences（5’-3’） | Usage*^a^* |
| --- | --- | --- |
| *ureA*-UP-F | CTCTATGGCAGTGGCTAC | a |
| *ureA*-UP-R | GTATCAGTCCAAACCATG |  |
| *ureA*-DN-F | TAATCAAAAAATTTCAGAAA | b |
| *ureA*-DN-R | CTCCTTAGGATCAATCCA |  |
| *ureB*-UP-F | CGCTGTCAAGGGCTTCAA | a |
| *ureB*-UP-R | CAGTTTTTGGAGCGACATG |  |
| *ureB*-DN-F | TAGATGAGCACAGTGATT | b |
| *ureB*-DN-R | TTCCTTCGAGCAATATTG |  |
| *ureC*-UP-F | CTGCCAGAGGGTTTCTAA | a |
| *ureC*-UP-R | CAAGAAGGAGTCCCCATG |  |
| *ureC*-DN-F | TAAAGACCATCTTGTTAAAAAT | b |
| *ureC*-DN-R | ACTTACCCACGGGTAGAG |  |
| *ureG*-UP-F | GTCAGGGCAGCCCAGGGT | a |
| *ureG*-UP-R | TGGAGTGAATCAAAAATG |  |
| *ureG*-DN_F | TAACGTAAAAAACAAAAT | b |
| *ureG*-DN-R | TCGAGTTGTGCTGTCGTC |  |
| *ureD*-UP-F | GAAAAAGTTATCACCGGA | a |
| *ureD*-UP-R | CAATTTCAGACAAAAATG |  |
| *ureD*-DN-F | TAACCCATCCCTAGCTCC | b |
| *ureD*-DN-R | GCCAATGAATGGGGTCTT |  |
| *ureE*-UP-F | TCTCTACCGTTATGAAAT | a |
| *ureE*-UP-R | CATCCGAGGTCACTGGTG |  |
| *ureE*-DN-F | TGACCCCCAGGGGTTAAG | b |
| *ureE*-DN-R | CCACGGTGGGCGTACCTA |  |
| *ureF*-UP-F | ATCCCAATGCATTGGCGG | a |
| *ureF*-UP-R | ATATCGCTAAAGACGATG |  |
| *ureF*-DN-F | TAAGCACACAATCTGGGC | b |
| *ureF*-DN-R | TGGGGATCACCTTTGCTG |  |
| *ureC*-F | ATACTCGCGCCTT | c |
| *ureC*-R | CAACGCTTTATTGGAAGATTGA |  |
| *UreG*-F | CCGGAATTCGTCAGGGCAGCCCAGGGT | c |
| *UreG*-R | TGCTCTAGAGACGACAGCACAACTCGA |  |
| *Synechococcus* sp. WH 8102 *-ureG*-F | GTGTGAGGAAAAACAAGGGCCCGGGTACCGTATTCCCGCTTGTTTCGGAG | c |
| *Synechococcus* sp. WH 8102 *-ureG*-R | TGTAAACAGTTTATTCATCTCGAGGTACCTCAAGCATTCGGTAGCTGTTGC |  |
| *Prochlorococcus* MIT 9313 *-ureG-*F | AGCTCTCCCATG | c |
| *Prochlorococcus* MIT 9313 *-ureG-*R | CCAAATCGCCCTTAG |  |
| *Helicobacter* sp *-ureG*-F | CGGGGTACCATACTCGCGCCTT | c |
| *Helicobacter* sp *-ureG*-R | CCCTCGAGTCAATCTTCCAATAAAGCGTTG |  |

*a* Letters have the following meanings: a, used to amplify corresponding gene homologous upper arm; b, used to amplify corresponding gene homologous arm downstream; c, used to amplify target genes

Table S2 The discharge of urea and nitrogen pollution source in selected ocean areas.

| Ocean Areas | Pollution sources | Reference | Year |
| --- | --- | --- | --- |
| global surface sea | / | Sipler & Bronk^9^ | 2015 |
| global surface sea | / | Robert T. Letscher^10^ | 2013 |
| Worldwide Coastal | Domestic sewage; Agricultural sewage | S. P. Seitzinger^11^ | 2010 |
| Northeast Brazil sea | Village Sewage； Domestic sewage | Daniela Mariano Lopes da Silva^12^ | 2018 |
| southwest Indian | Domestic sewage;  Agricultural sewage | Mintu Elezebath George^13^ | 2021 |
| Mediterranean Sea | urban effluents; industrial discharges; fertilizers | Michael Karydis^14^ | 2012 |
| southeastern Spanish sea | nitrogen fertilization | Miguel Lorite-Herrera ^15^ | 2008 |
| sub-Saharan Africa (SSA) | Agricultural sewage; industrial sewage; Domestic sewage | P.M. Nyenje^16^ | 2010 |
| Red Sea | Domestic sewage | David Peña-García^17^ | 2014 |
| Lima, Peru Peru Rimac river inlet | human feces and domestic sewage | Pedro E. Romero ^18^ | 2021 |
| Tanzania River Inlet | Urea source: fertilizers, herbicides, pesticides, excretion of animals | J. R. Selemani^19^ | 2017 |

Table S3 Strains used in this study.

| **Strain** | **Instruction** | **Source** |
| --- | --- | --- |
| *Synechococcus* sp. PCC 7002 | Wild type | Laboratory preservation |
| Mut-*ureA* | Km^R^, The *ureA* gene of *Synechococus* sp. PCC 7002 is mutated | This study |
| Mut-*ureB* | Km^R^, The *ureB* gene of *Synechococus* sp. PCC 7002 is mutated | This study |
| Mut-*ureG* | Km^R^, The *ureG* gene of *Synechococus* sp. PCC 7002 is mutated | This study |
| Mut-*ureC* | Km^R^, The *ureC* gene of *Synechococus* sp. PCC 7002 is mutated | This study |
| Mut-*ureD* | Km^R^, The *ureD* gene of *Synechococus* sp. PCC 7002 is mutated | This study |
| Mut-*ureE* | Km^R^, The *ureE* gene of *Synechococus* sp. PCC 7002 is mutated | This study |
| Mut-*ureF* | Km^R^, The *ureF* gene of *Synechococus* sp. PCC 7002 is mutated | This study |
| Com-*ureC* | Km^R^, Sp^R^. Complementation of the *ureC* gene of *Synechococcus* sp. PCC 7002 *ureC* mutant strain | This study |
| Com-*ureG* | Km^R^, Sp^R^. Complementation of the *ureG* gene of *Synechococcus* sp. PCC 7002 *ureC* mutant strain | This study |
| Com-WH 8102 | Km^r^, Sp^r^. The *ureG* of *Synechococcus* sp. WH 8102 was used to complement the *ureG* of *Synechococcus* sp. PCC 7002 *ureG* mutant strain | This study |
| Com-Pmt 9313 | Km^r^, Sp^r^, The *ureG* of *Prochlorococcus* MIT 9313 was used to complement the *ureG* of *Synechococcus* sp. PCC 7002 *ureG* mutant strain | This study |
| Com-Hp | Km^r^, Sp^r^, The *ureG* of *Helicobacter* sp was used to complement the *ureG* of *Synechococcus sp*. PCC 7002 *ureG* mutant strain | This study |
| **Eukaryotic algae** | **Instruction** | **Source** |
| *Thalassiosira pseudonana* | Wild type | Laboratory preservation |
| *Karenia mikimotoi* | Wild type | Laboratory preservation |
